# Quantum Theory on Genome Evolution

**DOI:** 10.1101/034710

**Authors:** LiaoFu Luo

## Abstract

A model of genome evolution is proposed. Based on several general assumptions the evolutionary theory of a genome is formulated. Both the deterministic classical equation and the stochastic quantum equation are proposed. The classical equation is written in a form of of second-order differential equations on nucleotide frequencies varying in time. It is proved that the evolutionary equation can be put in a form of the least action principle and the latter can be used for obtaining the quantum generalization of the evolutionary law. The wave equation and uncertainty relation for the quantum evolution are deduced logically. Two fundamental constants of time dimension, the quantization constant and the evolutionary inertia, are introduced for characterizing the genome evolution. During speciation the large-scale rapid change of nucleotide frequency makes the evolutionary inertia of the dynamical variables of the genome largely decreasing or losing. That leads to the occurrence of quantum phase of the evolution. The observed smooth/sudden evolution is interpreted by the alternating occurrence of the classical and quantum phases. In this theory the probability of new-species formation is calculable from the first-principle. To deep the discussions we consider avian genome evolution as an example. More concrete forms on the assumed potential in fundamental equations, namely the diversity and the environmental potential, are introduced. Through the numerical calculations we found that the existing experimental data on avian macroevolution are consistent with our theory. Particularly, the law of the rapid post-Cretaceous radiation of neoavian birds can be understood in the quantum theory. Finally, the present work shows the quantum law may be more general than thought, since it plays key roles not only in atomic physics, but also in genome evolution.

## Introduction

Genome is a well-defined system for studying evolution of species. There are many topics on genome evolution. A central problem is genome size evolution. The C-value enigma is still puzzling and perplexing [1][2]. How to understand the time arrow of the evolution? [3] The directionality means there possibly exists a set of basic equations for genome that evolves with respect to time *t*. The time *t* is a fundamental variable in the evolutionary theory. To find the genome evolutionary equation with respect to *t*, this is the first motivation of the present article. The evolution of species is a very complex process. The phylogenetic tree provides an appropriate quantitative description on evolution. However, although there have been proposed many mathematical methods to deduce evolutionary trees, it seems lacking a dynamical foundation for reconstruction of the tree from the genome sequence data. In my opinion, the reconstruction of evolutionary tree should be reconciled to the finding of the genome evolutionary equation. Then, what is the dynamical variable appropriate to reconstruct the evolutionary tree and deduce the genome evolutionary equation simultaneously? Since the k-mer non-alignment algorithm (KNA) is a good tool to obtain the evolutionary tree [4], we try to use the nucleotide (k-mer) frequencies as a set of dynamical variables of the differential equations describing evolution. To establish a mathematical theory on genome evolution that can be reconciled to evolutionary tree reconstruction, this is the second motivation of the present article. Thirdly, from the evolutionary phenomenology, two theoretical points, phyletic gradualism and punctuated equilibrium, were proposed to explain the macroevolution. It seems that both patterns are real facts observed in fossil evolution [5]. A deeper research question is what conditions lead to more gradual evolution and what conditions to punctuated evolution, and how to unify two patterns in a logically consistent theory. To give a unified mathematical theory on genome evolution, this is the third motivation of the present article. We shall suggest a set of second-order differential equations for genomic nucleotide frequencies which varies in time. Then, by putting the differential equation in a form of the least action principle, we shall demonstrate that the classical evolutionary trajectory can generally be replaced by the trajectory-transitions among them. Thus, the concept of quantum evolution is introduced. Both classical phase (gradually and continuously) and quantum phase (abruptly and stochastically) that have been observed in the evolution can be explained in a natural way. Fourthly, what rules are obeyed by the new species production? ‐‐ this is a difficult problem in existing evolutionary theory. However, the quantum theory gives us a fully new view on genome evolution. We shall demonstrate that the problem can be treated in a quantitative way, demonstrate that the speciation events obey a statistical law and the speciation probability can be calculated from the point of quantum transition. To give a formulation to study the new species production, this is the fourth motivation of the present article.

The extinction of the dinosaurs and pterosaurs at the end of Cretaceous (66 millions of years ago) left open a wide range of ecological niches, allowing birds and mammals to diversity. Most of the vacated avian niches were thought to have been filled by a rapid post-Cretaceous radiation of neognathous birds which today include over 99% of all bird species.[6]–[9]. Moreover, avian genomes show a remarkably high degree of evolutionary stasis at the level of chromosomal structure, gene syntency and nucleotide sequence. The macrochromosome studies reveals that the ancestral pattern has remained largely unaltered in the majority of avian genomes[10]. How to understand the law of the rapid post-Cretaceous radiation of birds, more than ten thousands of avian genomes producing explosively in a period shorter than 10^7^ years? How to explain the rapid accumulation of information in the early stage and soon afterwards the evolutionary stasis for avian genomes? To answer these questions we need a quantitative theory of evolution. On the other hand, the bird macroevolution was explored recently by using full genomes from 48 avian species representing major extant clades [11][12]. Then a more comprehensive phylogeny of 198 avian species was established by using targeted next-generation DNA sequencing [13]. The abundant data for avian species afford a splendid field for investigation of the genome evolution. In the present article we shall use the proposed quantum theory of evolution to study avian genome and explain the evolutionary data.

The k-mer frequency distribution is a characteristic of a genome (genome barcode) [14][15]. Several years ago we pointed out that the evolutionary tree can be deduced by non-alignment method – k-mer frequency statistics. By using 16S rRNA (18S rRNA) as molecular clock, we found that the phylogenetic trees deduced from k-mer frequency agree well with life tree when k≥7 [4]. If the genome growth is a fully stochastic process then DNA should be a random sequence and the k-mer frequency in the sequence takes Poisson distribution with the most probable value 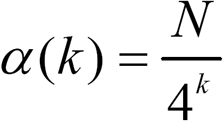 where *N* is genome length. However, the non-random force does exist in genome evolution. The most important non-random forces include functional selection and sequence duplication, etc. Due to the non-random forces the k-mer frequency deviates from Poisson distribution. Set 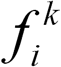 is the frequency of the *i*-th k-mer. The square deviation 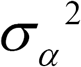 expressed by 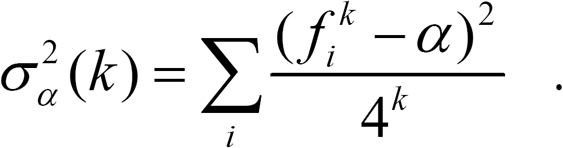. The ratio of the deviation 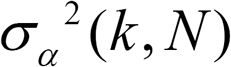 for a genome sequence of length *N* to the deviation *α*(*k, N*) for a random sequence of same length represents the nucleotide correlation strength or non-randomness level of the genome. It was found that there exists a meaningful relation between correlation strength and genome complexity [14]. Aiming to find the dynamical variable of evolution we used k-mer non-alignment algorithm (KNA) to reconstruct the evolutionary relationship for avian genomes.[16] Denote the probability of base *a* (*a* =A,G,C or T) occurring in a sequence by *P_a_*, and the joint probability of base *a* and *b* occurring sequentially in the sequence by *P_ab_*. In general, let *σ* = *abc*… being an oligonucleotide *k* bases long, we denote the joint probabilities of the bases in *σ* occurring in the sequence by *p_σ_*. For any given *k* we always have 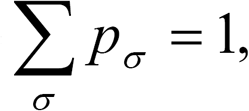 where the summation over *σ* is over the set {*σ*} of the 4^*k*^ oligonucleotides of length *k*. Given two sequences Σ and Σ^’^ with sets of joint probabilities {*p_σ_*} and {*p_σ_*’}, respectively, define a distance, called a *k*-distance, between the two sequences based on the difference of joint probabilities in the two sets as follows

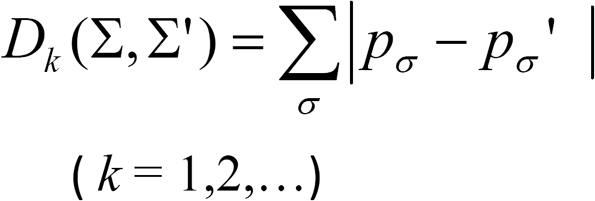

For long sequences a *k*-distance is insensitive to minor misalignments between two sequences. By using sequence data for 47 representative avian genomes we calculated the *k*-distances and deduced the phylogenetic tree for each *k*. We found that the quality of tree is gradually improved and the tree topology is stabilized with increasing *k* and reaches a plateau at *k*=11 or 12. The tree deduced by KNA algorithm (Figure A1 in Appendix) is consistent with experimental evolutionary relationship and basically in accordance with those obtained in literatures [12][13]. Above calculations show that the k-mer frequency is a good dynamical variable for describing genome evolution. This provides a conceptional basis for studying evolution through a set of evolutionary equations for nucleotide frequencies in the genome.

**Figure A1.**
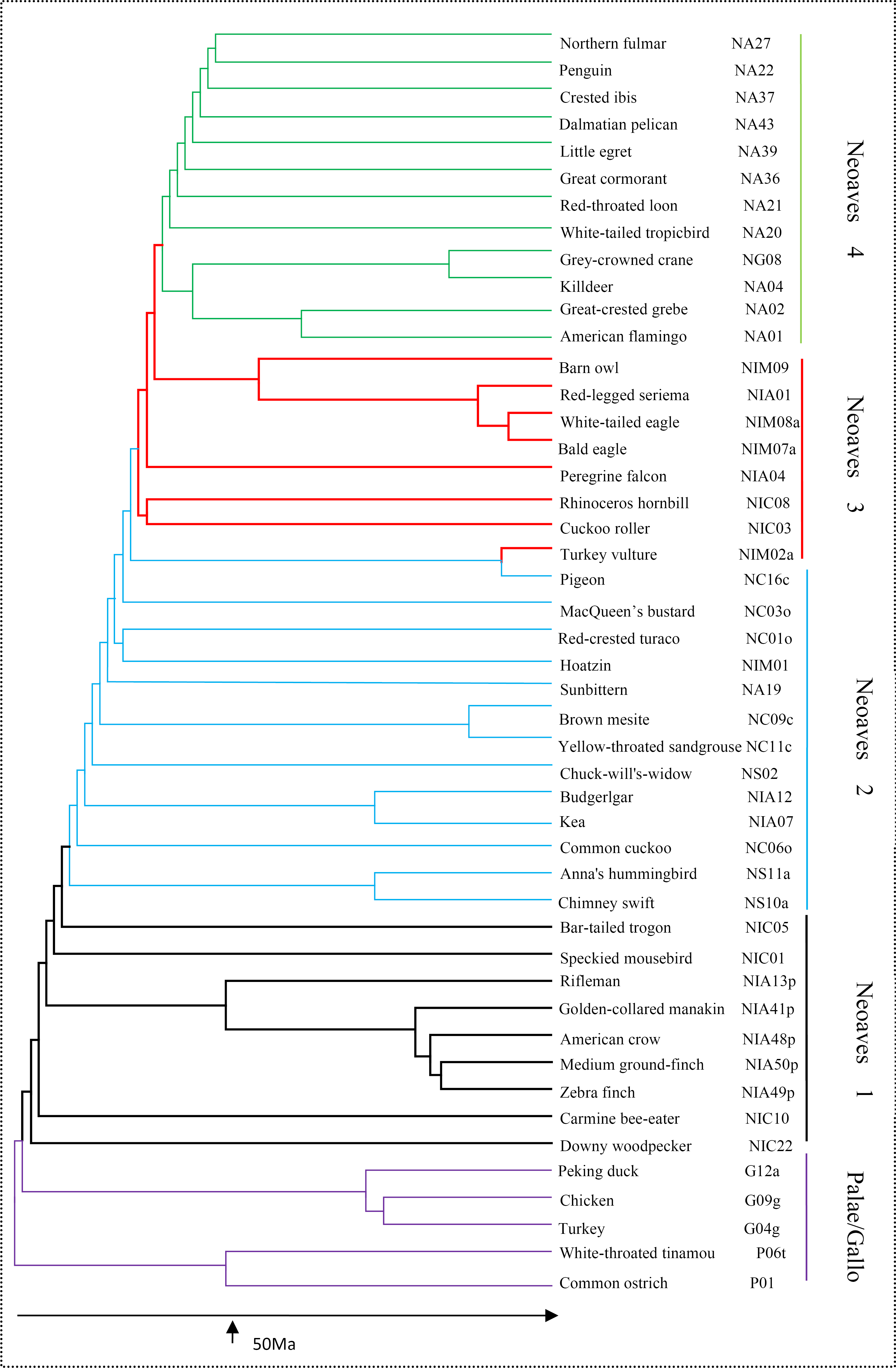
Phylogenetic tree deduced by KNA algorithm. The evolutionary tree of 47 birds was deduced in [16] that can be compared with trees published in [12] [13].

The materials are organized as follows. Firstly, three basic assumptions on the quantum theory of evolution are formulated. The classical evolutionary equation of the genome is written in terms of nucleotide frequencies varying with time. The evolution is described by a second-order differential equation. The path-integral quantization of the equation leads to the quantum transition between classical trajectories. To apply the theory to the practical system the supplementary assumptions on the form of evolutionary potential are proposed. In the subsequent sections the classical and the quantum motions of the genome are respectively discussed. In the classical phase evolution, the environmental potential parameters, evolutionary inertial parameter and dissipation parameter will be estimated by fitting avian genomic data. The rapid accumulation of information and the evolutionary stasis for avian genomes will be explained. In the quantum phase evolution the discreteness of quantum state and the ground-state wave function of avian genome are deduced. New species production are calculated by the quantum transition between discrete quantum states and based on the calculations the rapid post-Cretaceous radiation of neoavian birds can be understood more directly. In the final section, a concluding remark will be indicated that the quantum law may be more general than thought, since it plays key roles not only in atomic physics, but also in genome evolution.

## Basic assumptions of the quantum theory on genome evolution

**Ansatz 1**: For any genome there exists a potential *V*(*x*_1_,…,*x_m_,t*) to characterize the evolution where *x_i_* means the frequency of the *i*-th nucleotide (or nucleotide *k*-mer, *m* = 4^*k*^ in DNA.

Since *V*(*x*_1_(*t*),…, *x*_m_(*t*),*t*) ≠ *V*(*x*_1_(−t),…,*x*_*m*_(*x*_m_(*−t*),*−t*) in general, the potential can be used to describe the evolutionary direction.

**Ansatz 2**: The genome evolution equation reads as

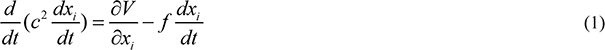

(1) where *f >*0 is a dissipation coefficient representing the effect of fluctuation force. The parameter
 *c*^2^ is introduced with the dimension of (time)^2^ which represents the evolutionary inertia of the dynamical variables {*x_i_*} of the genome.

The evolutionary law can be reformulated based on Feynman’s action integral. Introduce a functional (called information action)

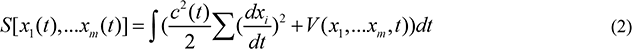

Then the solution of (1) can be expressed as

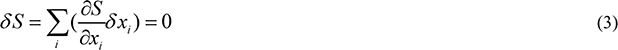

as the dissipation is weak(*f* can be neglected). Therefore, the classical evolutionary trajectory *x_i_*(*t*) satisfies the principle of the least action. By use of path integral quantization the evolutionary trajectory theory can be generalized to a more general quantum formalism.

**Ansatz 3**: The genome evolution obeys a general statistical law, described by the information propagator [17]

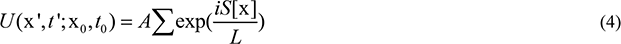

(x=(*x*_1_,…*x*_m_)). The summation is taken over all ideal paths satisfying x=x_0_ at *t*_0_ and x=x’ at *t*’ which is termed as functional integral in mathematics. (see Appendix 1)

Here *L* is a quantization constant of time dimension. The path integral *U*(x’,*t*’;x_0_,*t*_0_) describes the evolution of the genomic statistical state from *t*_0_ to *t*’. When the virtual variation δ*x*_i_(*t*) makes *δS≫L* all terms in the summation (4) will be canceled each other due to phase interference apart from those in the vicinity of classical trajectory where *S* takes a stationary value (*δS* =0). Therefore, the classical trajectory is a limiting case of the general quantum theory and the constant *L* can be looked as the threshold for classical approximation. Since during speciation the definite classical trajectory has lost its meaning and instead the quantum picture of trajectory transitions holds, we assume the quantization constant *L* is related to the time for new species formation. The constant *L* is species-dependent because of the generation time and therefore the speciation time varying largely from species to species.

**Supplementary assumptions** To make concrete analysis on genome evolution we further assume

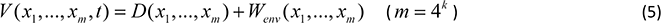

*W_env_* is a selective potential dependent of environment, and *V* depends on *t* through the change of environmental variables. *D* means the diversity-promoting potential

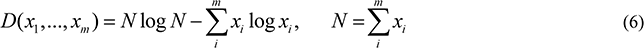

(hereafter the notation log means log_2_) Notice that the potential defined by (5) equals Shannon information quantity multiplied by *N*. Set 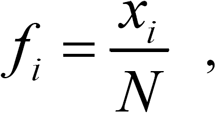 one has

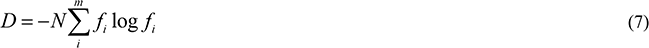

If the correlation between bases can be neglected then the diversity of *k*-mers is *k*-fold of the diversity of single nucleotides. In literatures *D* is called diversity measure which was firstly introduced by Laxton [18]. In their studies the geographical distribution of species (the absolute frequencies of the species in different locations) was used as a source of diversity. Recently, the method was developed and applied successfully to various bioinformatics problems, for example, the intron splice site recognition[19], the promoter and transcriptional starts recognition[20], the protein structural classification[21], the nucleosome positioning prediction[22], etc. Now we shall use it to study evolutionary problem. Many examples show that the genome always becomes as diverse as possible and expands their own dimensionality continuously in the long term of evolution [3]. These observations are in agreement with the evolution-promoting potential given by Eq (6). In her book “*Investigation*” Kauffman wrote：“Biospheres, as a secular trend, that is, over the long term, become as diverse as possible, literally expanding the diversity of what can happen next. In other words, biospheres expand their own dimensionality as rapidly, on average, as they can.” and called it “the fourth law of thermodynamics for self-constructing systems of autonomous agents” [23]. Our proposal on evolution-promoting force is consistent with Kauffman’s suggestion about the force for “expanding the diversity” in autonomous agents.

Assume the environmental potential *W*_*env*_ composed of two parts,

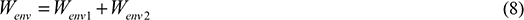

*W*_*env*1_ is only a function of 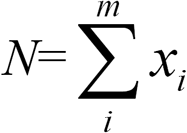 that is symmetrical relative to m oligomers and *W*_*env*2_ selective potential asymmetrical to m components. As we know that for the case of small kinetic energy 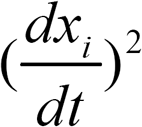 the classical trajectory of a mechanical system is always near the bottom of the potential energy. Through the mechanical simulation one may further assume the high-order term of *N* in *W*_*env*1_ is neglected. Thus we have

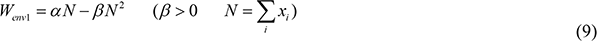

Note the form of *W*_*env*1_ is invariant under the transformation of the oligomer length from *k* to *k*-1. Different from *W*_*env*2_, which peaks at some particular sets of oligomers, the environmental selection of *W*_*env*1_ is only correlated with the total length N, irrespective of particular selection of oligomers. The selection of *W*_*env*1_ with *N* and called negative as *W*_*env*1_ decreases with *N*. As *β*>0 and 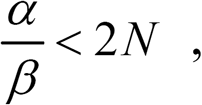, the environmental selection of *W*_*env*1_ is always negative. The parameters *α* and *β* in environmental potential are assumed to be constant corresponding to the stable evolution. However, *α* and *β* may change with *t* in the varying environment and the genome will select new evolutionary trajectory (new solution of Eq (1)) to adapt the environment. Sometimes, the sudden change of the environment makes *β*(*t*) increasing too rapidly and the genome may be shrunken and degenerate and the old species may be close to extinction.

## Results and discussions

### 1 Alternating occurrence of classical and quantum phases

Classical phase means the smooth evolution obeying the classical deterministic law, while the quantum phase means the sudden evolution obeying the quantum stochastic law. The present model of genome evolution predicts the alternating occurrence of both phases.

Many different estimates for the rate of evolution were made from the fossil records. As compiled by Gingerich[24][25] four hundred and nine such estimates were reported and they vary between 0 and 39 darwins in fossil lineage. Paleobiological studies indicated that species usually change more rapidly during, rather than between, speciation events. The smooth evolution always occurs between speciation events and the sudden evolution preferably occurs during speciation. The former can be interpreted as the classical phase and the latter as the quantum phase in our model. Paleobiological studies also indicated that the structurally more complex forms evolve faster than simpler forms and that some taxonomic groups evolve more rapidly than others. All these observations can be interpreted by the alternating occurrence of classical and quantum phases and discussed quantitatively based on evolutionary equations proposed above.

Phyletic gradualism states that evolution has a fairly constant rate and new species arise by the gradual transformation of ancestral species. While punctuated equilibrium argues that the fossil record does not show smooth evolutionary transitions. A common pattern is for a species to appear suddenly, to persist for a period, and then to go extinct. Punctuated equilibrium states that evolution is fast at times of splitting (speciation) and comes to a halt (stasis) between splits. The theory predicts that evolution will not occur except at times of speciation (Fig 1). It seems that phyletic gradualism and punctuated equilibrium are contrasting and contradicting theories.[26] However, from our model of genome evolution both smooth and sudden phase should occur in a unifying theory. By using the evolutionary equation (1) it is easily to deduce that the evolution has a range of rates, from sudden to smooth. Moreover, from the general formalism given by Eq (4) the evolutionary trajectories during speciation should be switched to quantum transitions among them. Thus, from the present theory the punctuated equilibrium and the phyletic gradualism are only the approximate description of two phases of an identical process.

**Figure 1.**
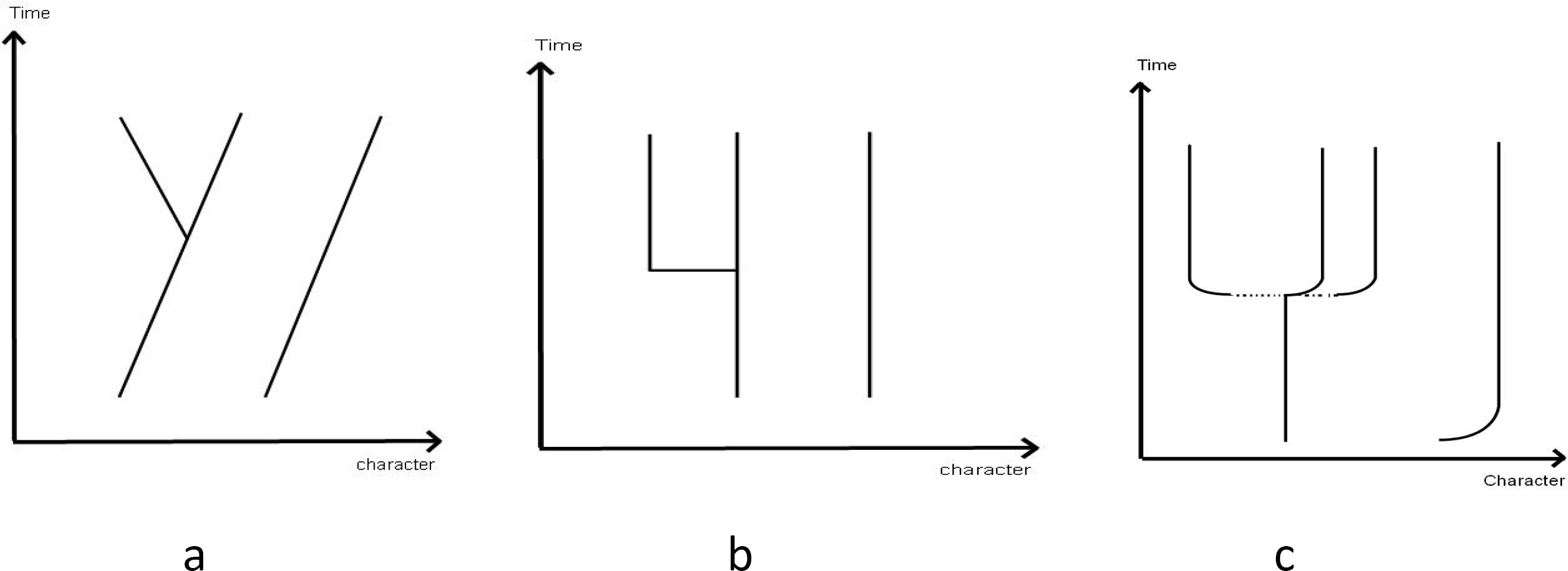
Comparison of three evolutionary theories. (a) phyletic gradualism, (b) punctuated equilibrium, (c) quantum evolution (dotted line means quantum transition)

### 2 Laws in classical phase

From (1) and (5)(6) the classical genome evolution equation reads as

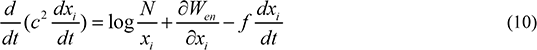

The first term of RHS of Eq (9) comes from the diversity potential which increases with 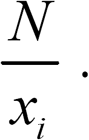. The less 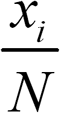 the stronger the force.

Consider single nucleotide evolution (*k*=1 case) first. The observed informational redundancy *R_I_*

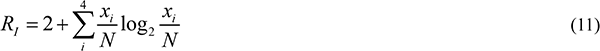

is small for most genomes except several prokaryotes that shows a small G+C content (for example, as small as 20% to 30% for some bacteria). Therefore, the asymmetrical potential *W*_*env*2_ can be neglected and one can approximate *W*_*env*_ to *W*_*env*1_ for most genomes. To obtain more detail understanding on classical evolutionary trajectory, we shall study avian genome as an example. From the full-genome sequence data of avian species [11]–[13] it is found the size *N*(sequence length) of genomes takes a value between (1044 – 1258)×10^6^ (Table A1 in Appendix 3). The informational redundancy *R_I_* takes a value between 0.008 – 0.026 that shows *x*_*i*_ very near to *N*/**4**.

Inserting Eq (9) into Eq (10) it leads to the equilibrium occurring at

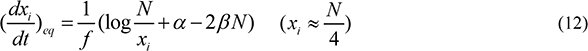

When *N* close to 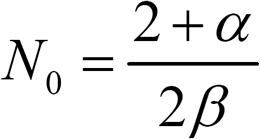 the genome size varies slowly and it tends to stabilization. We call it in stasis phase. Since the majority of avian genomes shows apparently the evolutionary stasis, if the parameter *α* and β are assumed same for different Aves then one may estimate β from the smallest length *N_small_* of the studied avian genomes,

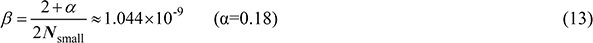

where α is assumed a small number between 0 and 2.

Through integration of Eq (10) one obtains

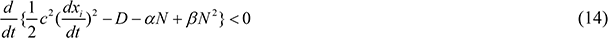

due to the damping effect of friction force. The dissipation of friction force is accumulated in time. In the initial evolution of a genome one may neglect *f* term in Eq (10) and deduce that *x*_*i*_ increases rapidly with a dimensionless acceleration

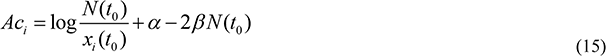

at *t=t*_0_.

Introducing dimensionless τ= *t/c* Eq (10) is rewritten as

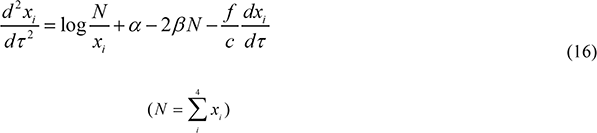

Eq (16) can be solved for each avian genome. The only parameter in equation (16) is *f/c* in addition to common environmental parameters α and β. The solutions for 12 Aves with branching time longer than 50 million years are shown in Fig 2. For example, the branching time of Common osterich is 51Mya and *N*= 1.228×10^9^ at present. In calculation the parameter *f/c*=0.0017 is chosen for Common osterich. We found x_*i*_ and *N* of this genome increase rapidly with the dimensionless time τ in a period of Δτ=3500 or the corresponding acceleration time Δ*t*=4M. Then, after the initial expansion the evlution comes to stasis. From Δ*t* and Δτ we obtain *c*=1142yr for Common osterich. The inertia of other genomes can be deduced in the same way.

**Fig 2.**
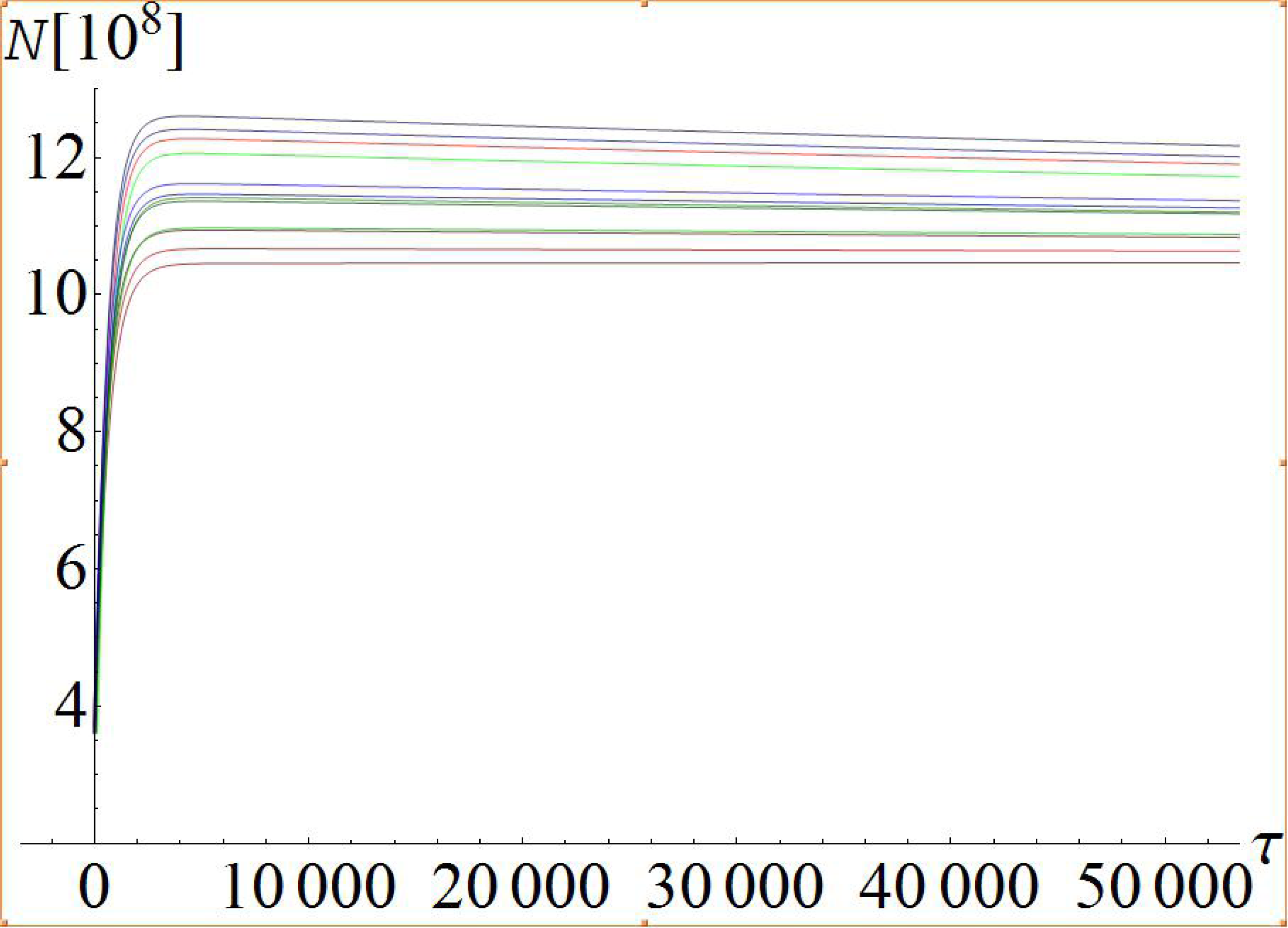
The evolutionary trajectory of 12 avian genomes^[32]^.

We found *c* obeys

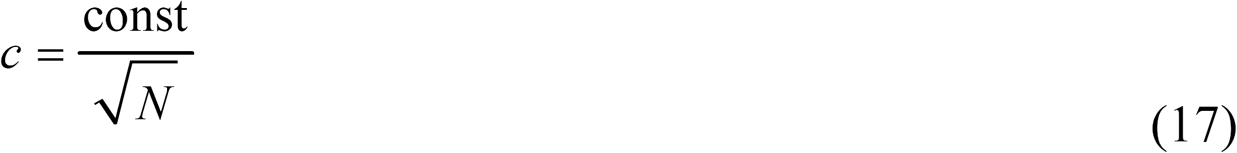

approximately. In fact, for 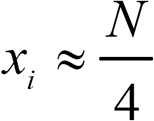 the equation (16) describes a damping oscillation whose frequency square oppositely proportional to sequence length. Considering that the acceleration time for all birds is nearly same, one can obtain Eq (17) immediately. On the other hand, from the input *f/c* and calculated *c* for Common osterich we obtain *f*=1.94 yr which is common for all Aves. Because many birds are in stasis phase from 50 Mya till now one estimates

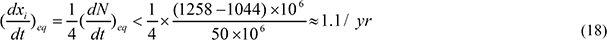

The dissipation force 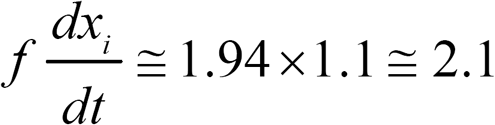 matches well with the promotion force 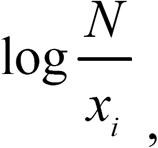 consistent with experimental data.

The genome length *N* versus the dimensionless time τ for 12 Ave (Penguin, Crested ibis, Common osterich, Hoatzin, White-tailed tropicbird, Cuckoo-roller, Red-throated loon, Red-crested turaco, Sunbittern, Mac Queen’s bustard, Carmine bee-eater, and Rifleman from top to bottom) are plotted. N increases rapidly and attains maximum at τ smaller than 4M. The parameters β = 1.0435×10^−9^, *f* = 1.94 yr are taken and *c* can be found in Table A1 in Appendix. The evolutionary inertia *c* increases for 12 genomes from top to bottom.

Note that in above calculation the environmental parameter is supposed to be a constant. However, if one assumes that β changes with time in the early stage of *N* expansion, the evolutionary trajectory will retain its basic character as shown in Fig 2 since *βN*^2^ term in equation is small as *N* not too large.

Now we will discuss the evolution of oligo-nucleotide frequency of a genome. The *k*-mer (*k*>1) frequency evolution is closely related to the single-nucleotide frequency evolution (*k*=1 case) of a genome. The Eqs (1) (5)-(8) still holds for *k*-mer case (*k*>1). From Eqs (6) and (7) one has *D*(*m* = 4^*k*^) = *kD*(*m* = 4) for independent sequence where *D*(*m* = 4) means diversity of single nucleotide. That is, if the correlation between neighboring bases is switched off then the *k*-mer diversity equals *k*-fold of the single nucleotide’s. If the consistency between the single nucleotide evolutionary equation and the *k*-mer equation in weak correlation is required then the environmental potential *W*_*env*1_ = χ(*N*) for k-mer should be equal k-fold of *W*_*env*1_ = *αN − βN*^2^ (Eq (9)) for single nucleotide evolution. On the other hand, the asymmetrical environmental potential *W*_*env*2_ cannot be neglected definitely in the oligo-nucleotide evolution. Therefore, one obtains a similar set of equations for *k*-mer frequency as Eq (16)

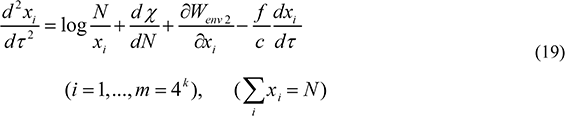

with

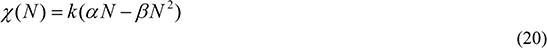

Here the environmental potential *W*_*env*2_ is responsible for the functional selection of some particular k-mers and can be written following the observed frequency distribution of k-mers in the studied genome.

### 3 Basic characteristics in quantum phase

The nucleotide frequency moving on classical trajectory and obeying deterministic equation cannot explain the new species production. The sudden change and the stochastic property in speciation event should be understood from a broad view. Suppose the statistical state of the DNA evolution is represented by a wave function *ψ*(x,*t*) that describes the probability amplitude (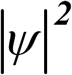 representing probability) of nucleotide frequency **x**(**x**={*x*_i_}) at time *t*. The propagation of wave function is determined by

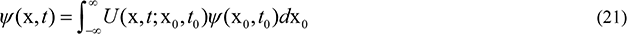

where *U* is given by Eqs (4) and (2). From Ansatz 3 we can prove *ψ*(x, *t*) satisfies Schrodinger equation (see Appendix 1)

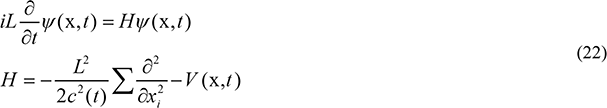

*H* is called Hamiltonian of the genome. The quantum evolutionary equation (22) is the logic generalization of the classical equation (1) (as *f*=0). The generalization is valid not only for the evolution in stable environment but also for the evolution in varying environment where the evolutionary potential *V* and inertia *c*^2^ are time-dependent. As a stochastically generalization of the deterministic equation the quantum evolutionary theory is applicable in studying the new species formation.

In addition to the least action principle and Feynman’s path integral approach the quantum generalization of the classic evolutionary equation can be deduced as well from Bohr’s correspondence principle. Niels Bohr indicated that there exists a good correspondence between classic-mechanical motion and quantum motion in atomic physics. As the mass of a particle is small enough the classical mechanics should be replaced by the quantum laws. The similar correspondence exists in present case. The inertia parameter *c*^2^ is time-dependent in general. During speciation the inertia of nucleotide frequency in the genome drops largely and the classical motion will be changed to quantum. Therefore, from both approaches we can deduce the stochastic evolutionary equation Eq (22), the quantum generalization of the deterministic equation (1). The deterministic equation describes the classical phase of evolution while the Schrodinger equation describes the quantum phase of evolution.

We will discuss the relations between classical and quantum evolutions in more detail. The time evolution of the genome is described by the propagation function 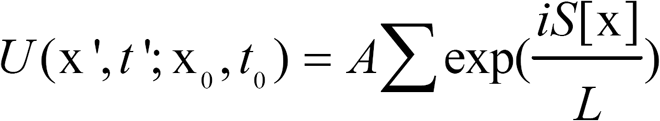 Assuming *L* is proportional to the average generation time λ for given species, one estimates *L=*(2000-3000) λ crudely from the human or bacterial speciation data. From Eq (2) the contribution of trajectory variation δx(*t*) to 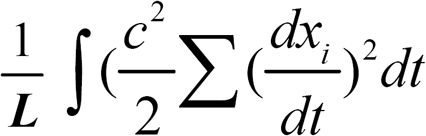 is proportional to *c*^2^. So the variation of phase *δS*/*L* in Eq (4) is much influenced by *c*^2^. The large *c*^2^ would make the contribution of different trajectories to the summation in (4), namely to *U*(x’,*t*’;x_0_,*t*_0_), easily canceled each other, while the small *c*^2^ does not. *c*^2^ is a constant for genome moving on classical trajectory under stable environment. However, during speciation the genome sequence undergoes tremendous changes and the evolutionary inertia of nucleotide frequency is lost, *c*^2^ jumping to a much lower value (*c*’)^2^. For example, one may assume *c*’ = 10^−3^*c* or less during speciation. The nucleotide changing rate 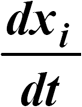 is assumed to be ν1/*T*. In the time duration *T* the variation of phase for different paths in functional integral

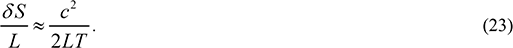

We have estimated *c*=1100yr for birds in previous section. Taking *L*=2000λ, λ=1yr and *T*=1yr one obtains *δS*/*L*=250. The phase variation is large enough and the strong interference makes only the classic trajectory retained. However, if the inertia is changed to (*c*’)^2^ = 10^−6^ *c*^2^ then *δS*/*L*= 0.00025 and the non-classical path cannot be neglected definitely. It means the evolutionary picture should be switched from classical to quantum and the quantum evolution occurs during speciation. To avoid confusion, in the following we shall use *c’* to denote evolutionary inertia in the quantum theory, instead of *c* on classic trajectory.

The uncertainty relation is a fundamental characteristic of quantum system. By using wave function ψ(x,***t***) one obtains the average of nucleotide frequency *x_i_*

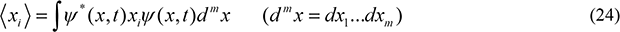

Define the conjugate of *x_i_* as 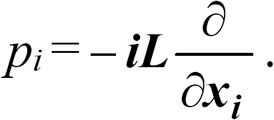 By using Schrodinger equation the average conjugate of frequency

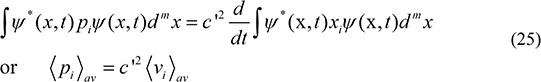

is deduced where *v_i_* means the changing rate of nucleotide frequency. Moreover, by defining square deviations

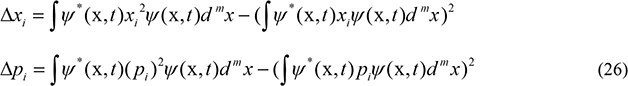

one easily obtains

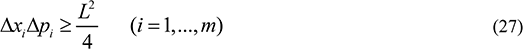

or

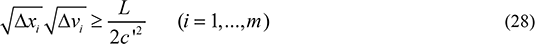

for any given time. Here 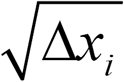 the uncertainty of nucleotide frequency and 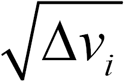 the uncertainty of the changing rate of nucleotide frequency. Eq (27) (28) means the uncertainty relation of nucleotide frequency and its changing rate in DNA sequence. Generally the RHS of inequality (28) is not negligible. Combining Eq (28) and Eq (23) we find the less the variation of phase the stronger the uncertainty. Since x and v cannot be determined simultaneously to an enough accuracy, the evolutionary trajectory cannot be accurately defined in principle. The small *c*’^2^ requires the simultaneous occurrence of the larger frequency deviation and the larger deviation of the frequency changing rate. This leads to the nucleotide frequency and the frequency changing rate no longer simultaneously measurable to an enough accuracy. Therefore, the picture of classical definite trajectory should be replaced by a large amount of rapid trajectory-transitions. Moreover, the uncertainty of nucleotide frequency 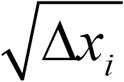 can be understood as the single nucleotide polymorphism. So Eq (28) gives a lower bound of the uncertainty for the changing rate of nucleotide frequency.

Now we shall discuss the quantum state of genome evolution. As an example, consider the avian genome evolution. Start from Eq (22) and assume *V*(x, *t*) given by (5)-(9). The quantum state described by the eigenstate of Hamiltonian is generally not continuously but discrete. To be definite we study the single-nucleotide frequency only. The eigenstate of *H* satisfies

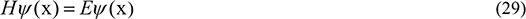

The Hamiltonian given by Eq (22) can be rewritten as

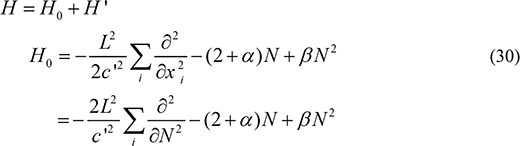

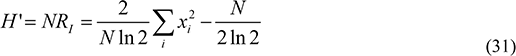

where Eqs (5)-(9) and（11）have been used and environmental potential *W*_*env*2_ neglected. *H*’ is a small quantity for avian genome (see the note after Eq (11)) which can be looked as a perturbation. Following perturbation theory, setting *ψ*(x) = *ψ*^(0)^(N)+*ψ*^(1)^(x), the zero order wave function *ψ*^(0)^ depends on *N* only，satisfying

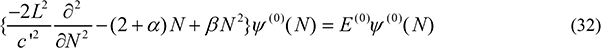

The parameters *α*, *β* and *c* take values at given time *t*_0_. The solution of Eq (32) is

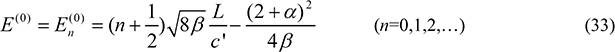

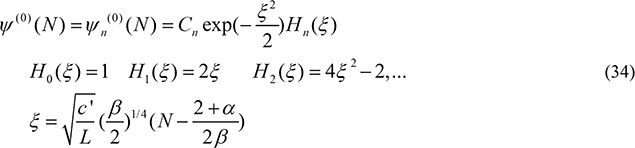

The first-order correction to 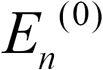 is

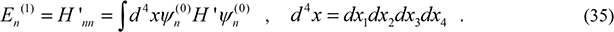

The first-order correction to wave function *ψ*^(0)^*N* can also be calculated from perturbation theory.

Eq (33) shows the Hamiltonian-level is equally spaced by 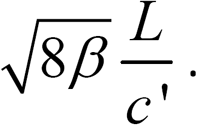 is replaced by the classical value *c*, estimated by 1100 yr for avian genome in previous section, then the spacing 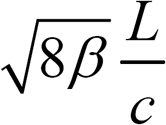 is a small quantity. So the eigenstates are basically continuous in classical phase of the evolution. However, during speciation the evolutionary inertia of nucleotide frequencies drops to a lower value as if in this time all evolutionary events happened more rapidly. If *c’* as a parameter of time dimension decreases to 10^−3^ or less of the classical value, *c*’ ≤ 10^−3^ *c*, then the picture of definite trajectory, namely *x_i_*(*t*) as a function of *t*, ceases to be correct and the state will be switched to quantum. A series of discrete eigenstates occur in quantum phase. In the quantum state the nucleotide frequency always takes some statistical distribution but not a definite value. Eq (34) shows the statistical distribution of *N* in ground state (*n*=0) peaks at 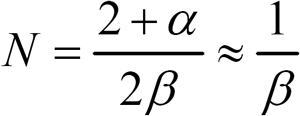 with width equal 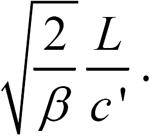.

Therefore, the quantum theory regards that the speciation event is essentially a quantum transition between initial “old” species and final “new” species. There always exists a Hamiltonian-level gap between low-lying ground state (*n*=0) and excited state (*n*=1,2,…), that can be seen from the large spacing 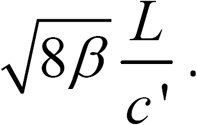 in a quantum genome. In fact, the gap of ground state should be deeper than 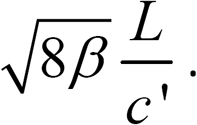 if the higher-order terms of *N*^3^ etc have been taken into account in the environmental potential equation (9) of *W*_*env*1_. Due to the deep gap occurring in the Hamiltonian ground level one should consider only the transition between ground states in studying speciation event.

### 4 Speciation studied from theory of quantum transition

In the following we shall discuss the speciation event from the view of quantum transition. Based on Schrodinger equation the speciation rate can be calculated. Suppose the initial wave function of the “old” species denoted by *ψ*_I_(x) satisfying Eqs (32)-(34) and the final wave function of the “new” species by *ψ*_F_(x) satisfying the same equation but *c*’ (*t*_0_), *β*(*t*_0_), *α*(*t*_0_)replaced by *c*’(*t_F_*), β(*t*_F_), α(*t*_F_) respectively. The transition from “old” to “new” is caused by a time-dependent interaction *H*_int_(*t*) in the framework of quantum mechanics. One may assume *H*_int_ comes from the time variation of evolutionary inertia and environmental potential, namely

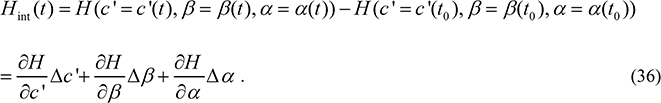

Thus the transitional probability amplitude is expressed by [27]

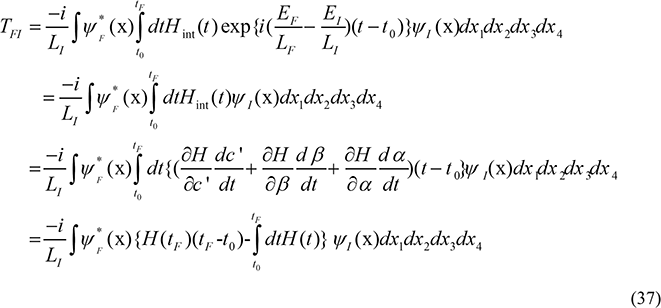

where

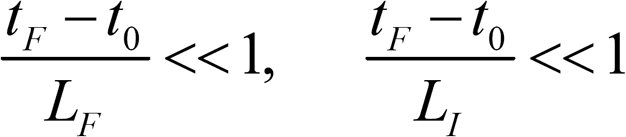

(*L_I_* and *L_F_* are the quantization constant of initial and final genomes respectively) has been taken into account and the partial integration has been used. On the other hand,

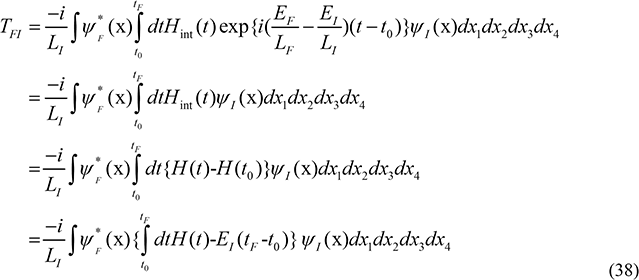

So,

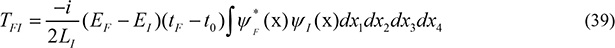

*E_I_* and *E_F_* – the eigenvalue of *H*, given by Eq (33) with *n*=0 where parameters *c*', *β*, *α* take values at *t*_0_ and *t_F_* respectively. The transitional probability

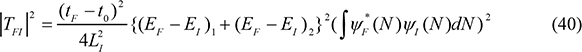

is determined by three factors, namely

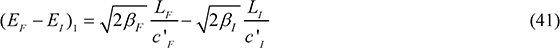

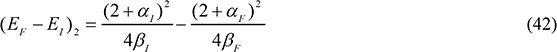

and the wave function integral 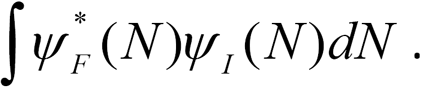 By using

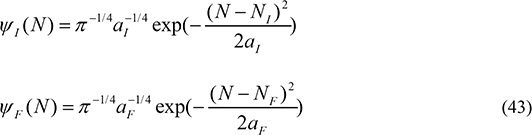

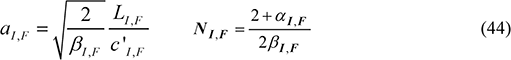

(*a_I, F_* – the frequency distribution width, and *N_I, F_* the frequency distribution centers for two genomes respectively, that can be found in Eq (34), the overlap integral can be calculated and it gives

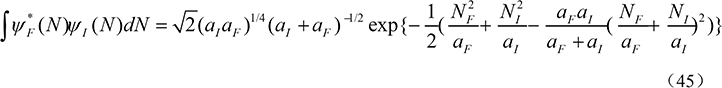

The transition time *t*_F_-*t*_0_ can be obtained by the normalization of the total transition probability from initial (*I*) to all possible final (*F*) states, namely

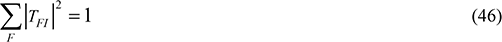

Under stable environment, *β_I_ = β_F_* and *α_I_ =ɑ_F_* one has (*E_F_ – E_I_*)_2_=0. The transition is determined only by (*E_F_ – E_I_*)_1_ and the overlap integral. However, under varying environment, considering 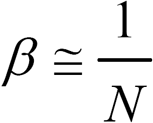 is a small quantity one has (*E_F_ – E_I_*)_1_<<(*E_F_ – E_I_*)_2_ generally. This explains why new species formation occurs most probably in the condition of varying environment. For example, a long-term experiment with *Escherichia coli* was made where the population evolved with glucose as a limiting nutrient. There was only a hypermutable phenotype appeared by about generation 26500.[28] However, in our experiments with *Escherichia coli* by repetitious ion irradiation, the colony polymorphisms of mutants were observed in the very early stage. Therefore, ion irradiation as a varying environment can effectively accelerate the genome evolution of *E. coli*.[29]. No loss of generality, in the following calculation we assume

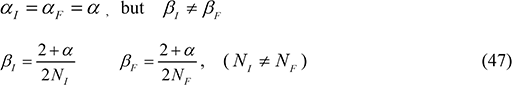

On the other hand the exponent factor in Eq (45)

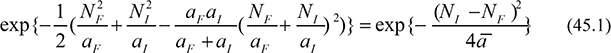

(*ā* is some average of *a_F_* and *a_I_* that shows the wave function integral or the transition is not negligible only when *N_F_* near *N_I_* which means *β_F_* near *β_I_*, the variation of environmental parameter being small during quantum transition. The point is understandable because *t_F_* – *t*_0_ is a small quantity and the changing environment only causes a small variation of environmental parameter β in such a short time. Moreover, Eq. (45.1) shows the transition probability is irrespective of the symbol of *N_I_* − *N_F_*. The sequence length of posterity genome can be longer, or shorter as well, than ancestry in quantum transition.

If the quantum transition occurs in stable environment, assuming 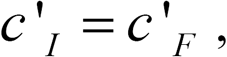, from (41) and (42) one has

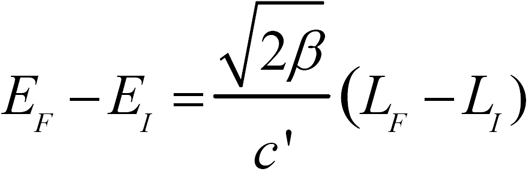

Set *L* proportional to the generation time λ. One obtains

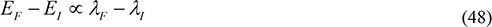

For several posterity genomes produced from one ancestry the branching ratio of transitional probability between different channels is proportional to (*λ_F_* − *λ_I_*)^2^ if the difference between the overlap integrals can be neglected. However, the speciation event often happens in varying environment and in this case the initial and final environmental parameters are different and one should use the original equation (40) to calculate the transitional rate.

The rapid post-Cretaceous radiation of neoavian birds provides a vast amount of experimental data to test the above quantum theory on new species production. Although different evolutionary trees ‐‐ for example, trees published in Science [12], in Nature [13] and IMU tree[16] ‐‐ show the basically same sister relationships among clades the earliest branches in Neoaves during the rapid radiation after the Cretaceous-Paleogene mass extinction event have quite different topology. We study 12 kinds of birds whose branching time was between 66 Mya to 50 Mya, namely Common ostrich, Carmine bee-eater, Rifleman, Sunbittern, Hoatzin, Mac Queen's bustard, Cuckoo-roller, Barn owl, White-tailed tropicbird, Red-throated loon, Crested ibis and Penguin (denoted as bird 1 to bird 12 respectively) and assume that they were produced from a quantum speciation event from one ancestry bird (denoted as Anc) at time *t* between *t*_0_ and *t*_F_. Then, these genomes of posterity birds follow their classical trajectories respectively after the quantum transition (Fig 2). The ancestry bird Anc is assumed to have the same characteristics, inertia c' and generation time λ, as the posterity bird 1, Common ostrich. The peak of the Anc genome sequence length is supposed at *pN*_1_ (*p* is a fraction 1/4,1/3 or 1/2 etc, *N*_1_ is the present genome length of Aves 1). The peak of the *i*-th posterity genome at time of quantum transition is supposed at (referred to Fig 2) [32]

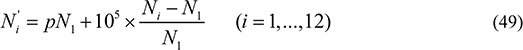

Where *N_i_* means the present sequence length of the *i*-th genome. By using Eqs (40)-(47) we calculate the transitional probabilities of new avian production and the results are summarized in Table 1.

**Table 1.** New avian production probability from ancestry Anc

| p | Tm(yr) | Aves1 | Aves 2 | Aves 3 | Aves 4 | Aves 5 | Aves 6 | Aves 7 | Aves 8 | Aves 9 | Ave10 | Ave11 | Ave12 |
| --- | --- | --- | --- | --- | --- | --- | --- | --- | --- | --- | --- | --- | --- |
| 1/4 | 0.88 | 0.1069 | 0.0459 | 0.0375 | 0.0731 | 0.1038 | 0.0718 | 0.0863 | 0.0745 | 0.1003 | 0.0903 | 0.1044 | 0.1051 |
| 1/2 | 0.85 | 0.1015 | 0.0533 | 0.0462 | 0.0770 | 0.0989 | 0.0761 | 0.0857 | 0.0752 | 0.0971 | 0.0895 | 0.0993 | 0.1003 |

Tm means the transition time *t*_F_-*t*_0_. Transition probabilities for 12 posterity Aves are listed in the table. In calculation c'=c/1000 and L=3000λ are assumed. The environmental parameter α=0.18 is taken, βI andβF are calculated from eq (47) where NI equal pN_1_ and N_F_ equal the average sequence length 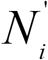 of the posterity Aves. (p=1/4 corresponds to classical trajectory given in Fig 2; for reference the case of p=1/2 is also calculated and listed).

Although the rapid post-Cretaceous radiation of neoavian birds arouses many difficulties in reconstructing avian evolutionary tree we found the quantum evolution gives new insight to the understanding of the complexity in the problem. Moreover, several general laws on new species production can be deduced from the example of Aves genomes:

1. In the time of post-Cretaceous radiation because of the extensive horizontal gene transfer and the wide-ranging hybridization of avian genomes the genomic sequences of many birds undergo great changes and the classical evolutionary inertia *c*^2^ of nucleotide frequency in these genomes has lost. That is, the parameter *c*^2^ should be replaced by a much smaller (*c’*)^2^ and the classical motion will be changed to quantum. In quantum evolution the speciation is a stochastic event, obeying a statistical law. Generally an ancestry produces several posterities with different probabilities.
2. There exist several discrete states for a genome and each state is characterized by an eigenvalue *E*. The difference of *E* between initial (ancestry) and final (posterity) genomes, ***E_F_ − E_I_***, is an important factor to determine the transitional probability between them.
3. In the quantum state of a genome, the sequence lengths obey a statistical distribution described by some wave functions. The overlap integral of the ancestry and the posterity wave functions is another factor to determine the transitional probability. The transition rate is not too small only when the peaks of two distributions in initial and final states are near to each other.
4. The transition time *t_F_* − *t*_0_ is very short as compared with the parameter *L*, 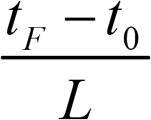 being about 1/1000 for birds as seen from Table 1. The above calculation remains valid, irrespective of different choices of parameters *L* and *c’* (for example, *L* changing from 3000 λ to 2000 λ and *c'* decreasing to a value much smaller than *c*/1000). It means the quantum transition of speciation event occurs abruptly and stochastically in one or few generations. The result of very short transition time is consistent with the concept of quantum transition in a general quantum theory.
5. The quantum transition preferably occurs in varying environment, the environmental parameter *β_I_* ≠ *β_F_*. The severe change of environment is beneficial for producing a new species. This provides an effective approach to search for the new species production. Note that due to the transition completed in a short time, the observable change of environmental parameter *β* during speciation may still be small even for severe environmental change.

## Conclusions

It is interesting to make comparison between the present quantum model for genome and the conventional quantum theory for electrons, atoms and molecules (even for biological macromolecules [30]). Both systems have wave function satisfying Schrodinger equation. Both theories have their classical limit. The classical trajectory of an atom or an electron is given by the position coordinate of the particle as a function of *t*, while the classical trajectory of a genome is given by the nucleotide frequency of DNA as a function of *t*. The former is constrained by particle’s energy while the latter is constrained by genome’s information. Energy is conserved with time but information always grows in evolution. The atomic quantum theory has an elementary constant, the Planck’s constant, while the genomic quantum theory has a corresponding quantization constant *L*. The Planck’s constant is universal and has dimension of energy but the genome constant *L* is species-dependent and has dimension of time. The classical limit of both theories can be deduced from Planck constant *h* or genome constant *L* approaching to zero respectively. However, the classical limit of atomic quantum theory is a new mechanics for macroscopic particle of large mass. While the genome inertia *c* is changeable in the course of evolution. For the same genome it changes thousand-fold in different phases of evolution. Two constants, quantization constant *L* and evolutionary inertia *c*, both in the dimension of time, play important roles in the present quantum evolutionary theory. The former is related to the realization of the quantum picture of the evolution, while the variation of the latter gives rise to the switch from classical phase to quantum phase or *vice versa*.

## Acknowledgement

The author is indebted to Dr Qi Wu for numerous helpful discussions. He also thanks Drs Liron Zhang, Xun Zhou, Yongfen Zhang and Guoqing Liu for their contributions in avian genome analyses and Dr Yulai Bao for his kind help in literature searching.

## Appendix 1 Deduction of wave equation in quantum phase

Suppose the statistical state of X=(*x*_1_,…*x*_*m*_ represented by a wave function **ψ**(x, *t*).

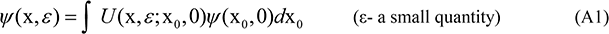

where *U* is a functional integral, expressed as

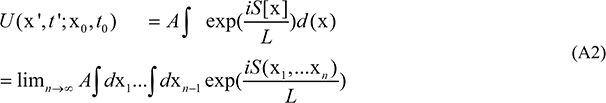

and *t*'−*t*_0_ = ε has assumed. Inserting the information action functional *S*[x] (Eq (5))

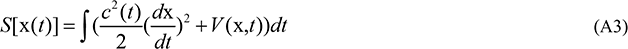

into (A2), after integrating over (x_1_,…x_*n*−1_) and taking limit *n* → ∞ we obtain [10]

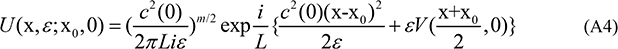

Eq (A4) has been normalized to *U* → *δ*(x-x_0_) as *V*=0. In the deduction of (A4) the integral formula

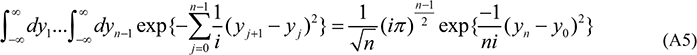

has been used. By using (A4), Eq (A1) can be written as

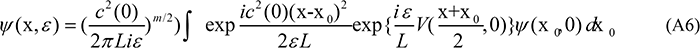

Here 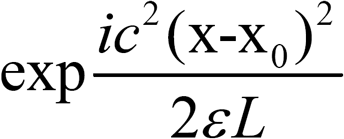 is a rapidly oscillating factor, only important near x-x_0_ =0. By setting x_0_ −x = *η*, Eq (A6) is rewritten as

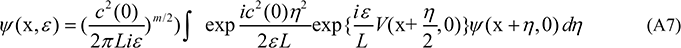

After expanding *ψ*(x + *η*, 0) and 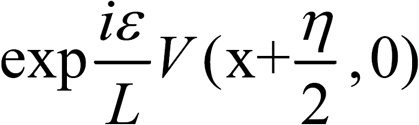 with respect to *η* and *ε* finally we obtain

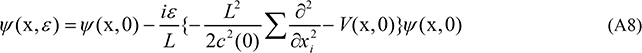

Therefore, *ψ*(x, *t*) satisfies Schrodinger equation,

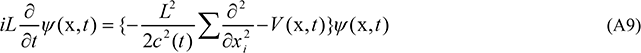

## Appendix 2 Phylogenetic tree of 47 representative birds

## Appendix 3 Notations and parameters of 47 Aves

**Table A1.**
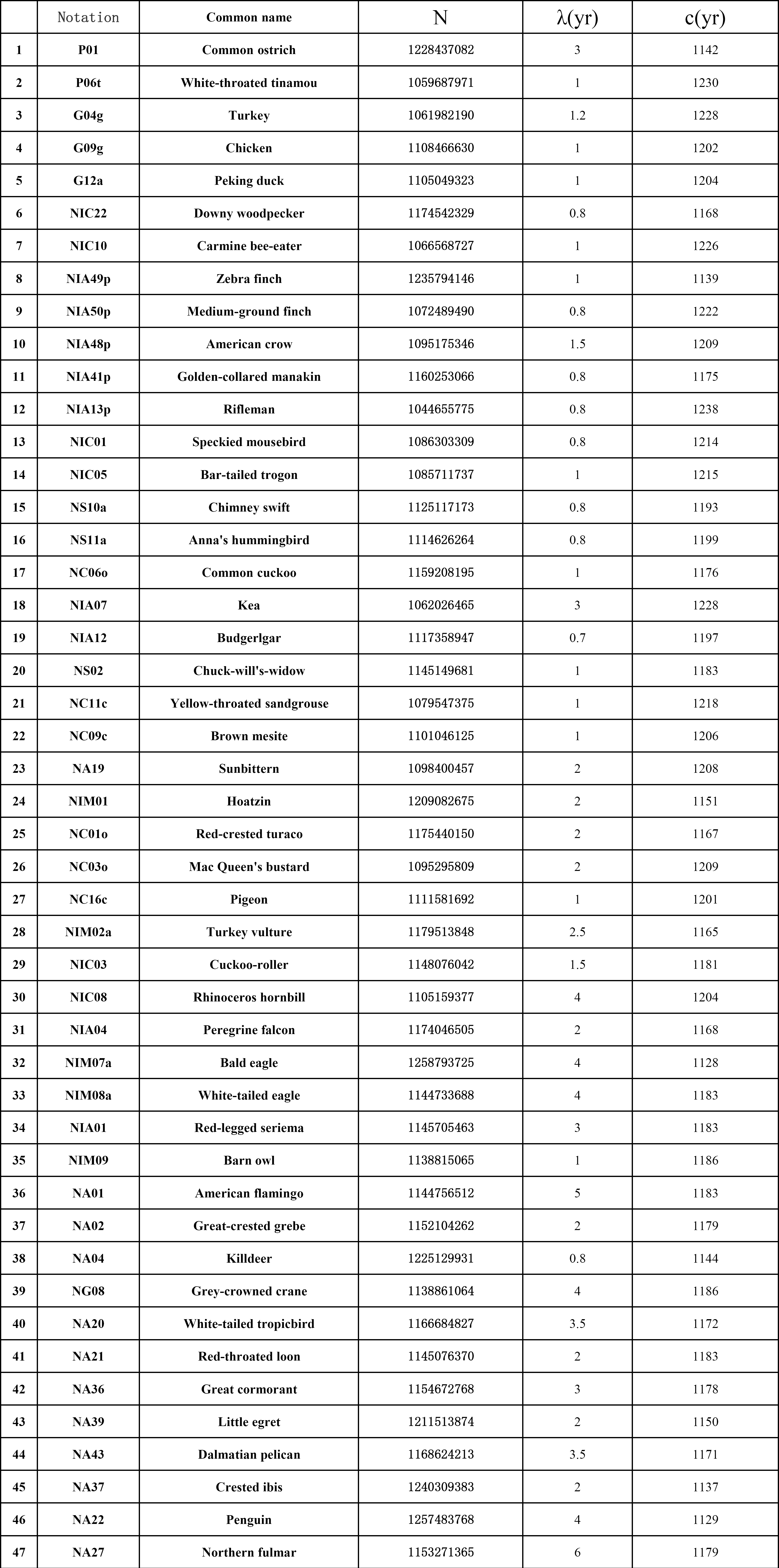
Parameters of 47 Aves used in theory of genome evolution

The notations of 47 birds are given by the following rules. Referred to 198 birds in ref [13], several letters are introduced to indicate the taxon of each bird. P01-P09 referred to Palaeognathae, G01-G16 referred to Galloanserae, NS01-NS13 referred to Neoaves-Strisores, NC01-NC16 referred to Neoaves-Columbaves, NG01-NG09 referred to Neoaves-Gruiforms, NA01-NA43 referred to Neoaves-Aequorlitornithes, NIM01-NIM10 referred to Neoaves-Inopinaves-Accipitriformes and related, NIC01-NIC26 referred to Neoaves-Inopinaves-Coraciimorphae, NIA01-NIA56 referred to Neoaves-Inopinaves-Ausralaves. The last lower-case letter indicates a subclass, for example, the letter p in NIA13p-NIA56p means the subclass passeriformes. These notations are also used in Fig A1. N means the genome sequence length. **λ** means the generation time, taking from [31]. c means the evolutionary inertia, calculated from the analysis of the classical trajectory of the genome evolution, see text.

